# Automated methodology for optimal selection of minimum electrode subsets for accurate EEG source estimation based on Genetic Algorithm optimization

**DOI:** 10.1101/2021.11.24.469917

**Authors:** A. Soler, L. Moctezuma, E. Giraldo, M. Molinas

## Abstract

High-density Electroencephalography (HD-EEG) has been proven to be the most accurate option to estimate the neural activity inside the brain. Multiple studies report the effect of electrode number on source localization for specific sources and specific electrode configurations. The electrodes for these configurations have been manually selected to uniformly cover the entire head, going from 32 to 128 electrodes, but electrodes were not selected according to their contribution to accuracy. In this work, an optimization-based study is proposed to determine the minimum number of electrodes and identify optimal combinations of electrodes that can retain the localization accuracy of HD-EEG reconstructions. This optimization approach incorporates scalp landmark positions of widely used EEG montages. In this way, a systematic search for the minimum electrode subset is performed for single and multiple source localization problems. The Non-dominated Sorting Genetic Algorithm II (NSGA-II) combined with source reconstruction methods is used to formulate a multi-objective optimization problem that concurrently minimizes 1) the localization error for each source and 2) the number of required EEG electrodes. The method can be used for evaluating the source localization quality of low-density EEG systems (e.g. consumer-grade wearable EEG). We performed an evaluation over synthetic and real EEG dataset with known ground-truth. The experimental results show that optimal subsets with 6 electrodes can obtain an equal or better accuracy than HD-EEG (with more than 200 channels) for a single source case. This happened when reconstructing a particular brain activity in more than 88% of the cases (in synthetic signals) and 63% (in real signals), and in more than 88% and 73% of the cases when considering optimal combinations with 8 channels. For a multiple-source case of three sources (only with synthetic signals), it was found that optimized combinations of 8, 12 and 16 electrodes attained an equal or better accuracy than HD-EEG with 231 electrodes in at least 58%, 76%, and 82% of the cases respectively. Additionally, for such electrode numbers, a lower mean error and standard deviation than with 231 electrodes were obtained.

**Highlights:**

- The number of EEG electrodes and their locations can be optimized for reconstructing the brain source activity.
- Optimally selected EEG electrodes can retain the accuracy of high density montages (256, 128 chs) for brain source estimation, when electrodes are selected according to their contribution to accuracy.
- With optimization, selected combinations of EEG electrodes will flexibilize the estimation of the source activity.

## 1. Introduction

Since the emergence of the international 10-20 system of electrode placement in 1958 (Jasper (1958)), electroencephalography (EEG) systems have gradually evolved towards high-density EEG (HD-EEG), first with the 10-10 system (Chatrian et al. (1988b,a)), then, with the geodesic electrode distribution to increase the number up to 256 channels (Tucker (1993)), and ultimately with the 10-5 system (Oostenveld and Praamstra (2001)) which sets the maximum number of EEG electrode positions with 345 locations for scalp coverage. While the 10-10 system was accepted as a standard by the American Clinical Neurophysiology Society and the International Federation of Clinical Neurophysiology (Sinha et al. (2016); Seeck et al. (2017)), the 10-5 system was not accepted by either of them (Acharya et al.(2016)). The 10-10 system and the 10-5 system were developed as extensions of the 10-20 system contemporaneously with the development of EEG source localization techniques which required a higher density of electrode settings to reduce localization error.

Accurate EEG source localization depends on many factors among which the density and location of scalp electrodes is an important one. It is widely accepted that an increased density of EEG electrodes generally provides more precise localization (Brodbeck et al. (2011); Lantz et al. (2003); Song et al. (2015)). Today, 128 EEG channel systems and even 256 EEG channel systems are common commercial choices (Suarez et al. (2000); Oostenveld and Praamstra (2001)). Even though the use of HD-EEG systems has resulted in improved spatial resolution (Pascual-Marqui (2002)) in source localization, practical use has come at a cost (Acharya et al. (2016); Jurcak et al. (2007)).

In Jurcak et al. (2007) an extensive investigation of the validity of the 10-20, 10-10 and 10-5 systems as relative head-surface-based positioning systems is presented. The authors argue that even though the 10-20 system has not been conceived to support localization of brain sources, its high-density extensions into the 10-10 and 10-5 systems have mainly provided increased electrode density proven to be effective in brain source localization (Pascual-Marqui (2002)). The high-density extensions of the 10-20 system logically inherit its electrode positioning principle, which was not conceived to improve the accuracy of brain source localization algorithms. Although it is sufficiently proven that these extensions are effective in increasing the accuracy of brain source localization, it remains to examine to what extent these high-density systems require all the electrode positions for attaining that. Sohrabpour et al. (2015) for example, concluded that increasing electrode number decreases localization error but this improvements plateaus at some point. These authors examined the relationship between localization error and the number of electrodes for the particular case of partial epilepsy in pediatric patients. The same is true however for every source/es being examined. In this regard, the questions we attempt to answer in this paper are: what is the minimum number of electrodes required for an accurate source localization of a particular brain activity? and, may it be possible to identify optimal low-density subsets of electrodes that can maintain the reconstruction quality of HD-EEG?

We propose an automated method that uses information from electrode locations on multiple systems and electrode configuration (e.g. the systems inherent to the 10-5, 10-10, 10-20 and/or geodesics systems) to select the optimal number of electrodes and their locations to solve the problem of EEG source localization for single and multiple sources. The optimization is based on the non-dominated sorting genetic algorithm II (NSGA-II) (Deb et al. (2002)). The algorithm combined with EEG inversion methods, searches for combinations of electrodes for solving the EEG inverse problem that minimizes the localization error while using the lowest number of channels possible. Considering *C* as the number of channels, exploring all the combinations of electrodes in order to find the optimal solution means solving the inverse problem 2^*C*^ times for single source case, and *S*(2^*C*^) for *S* multiple sources. These numbers will exponentially grow when increasing the number of channels, increasing significantly the computational efforts, e.g. in the case of 128 electrodes, it is required to solve the inverse problem 3.4*x*10^38^ times to evaluate all the possible electrode combinations. In contrast, the NSGA-II aims to reduce the computational cost in average to *O* ∗ (*P*^2^), where *O* is the number of objectives, in this case *s* + 1, and *P* the population size (Deb et al. (2002)). NSGA algorithms have been successfully applied in multiple fields for optimization and feature selection, like face expression recognition (Stoyell et al. (2021)), telecommunications (Huang et al. (2010)). This algorithm has been applied to EEG channel selection for classification of motor imagery Kee et al. (2015). Moreover, the algorithm has proven to be effective in identifying low-density EEG subsets that maximize classification accuracy while reducing the number of EEG channels required for epileptic seizure classification (Moctezuma and Molinas (2020a)) and subject identification (Moctezuma and Molinas (2020b)).

Previous works have evaluated sparse arrays of electrodes for source reconstruction. For example in (Jatoi and Kamel (2018), the authors have performed source reconstruction on real and synthetic EEG signals by using seven channels, where localization errors between 12 to 16 mm were found by using Multiple Sparse Priors (MSP) as inverse solver, however, no information about the electrode selection criteria was mentioned. In the work of Soler et al. (2020b), pre-preprocessing with Multivariate Empirical Mode Decomposition (MEMD) was successfully combined with MSP to improve the source reconstruction with low-density electrodes arrays of 32, 16 and 8 channels on real and synthetic EEG signals. In that study, the sub-set of electrodes were selected based on spatial coverage of the brain, but not optimized according to the source activity (in this paper not only MSP was reported, other inverse methods were also). In Soler et al. (2020a) a methodology based on regions of interest was proposed, where relevance analysis and electrode distance to the regions were the criteria for selecting subset of electrodes when solving the EEG inverse problem. The low-density subsets were combined with MSP and standardized low-resolution tomography sLORETA, obtaining localization errors below 10mm by using the 4, 8 and 16 most relevant electrodes from a high-density set of 60 channels. The error levels are remarkable and comparable to high-density levels. However, the study was limited by the evaluation on only synthetic EEG signals.

We approach the question of electrode reduction in a step by step manner, by first formulating a computer simulation-based source reconstruction problem for one single source and for multiple sources. In a second phase, we examine how effective is this automatic methodology in accurately detecting the stimulation site from intra-cerebral stereotactically implanted electrodes, based on the dataset recorded by Mikulan et al. (2020). For our study we combined three widely used source reconstruction algorithms, weighted minimum norm estimation (wMNE), sLORETA, and MSP, with the NSGA-II algorithm, with a particular emphasis on studying the effect of optimally reducing the number of electrodes, and investigating their locations for a specific source activity.

## 2. Materials and Methods

### 2.1. Methodology for channel selection based on genetic algorithm multi-objective optimization

EEG source reconstruction refers to the estimation of the properties of the source activity (source location, direction, and waveform) from the information registered by the electrodes on the scalp. For estimating this, the EEG inverse problem must be solved (The inverse problem is presented in more detail in section 2.2). The proposed methodology combines a source reconstruction algorithm and NSGA-II (Deb et al. (2002)), where the algorithm objective is to find electrode subsets with the minimum number of channels that retains the highest possible source localization accuracy (the accuracy of the reference HD EEG). For the optimization several inputs are required: The EEG or the event related potentials (ERPs) of the source(s) to analyze, the head model (this is required for source reconstruction, see section 2.2) and the ground-truth location of the source activity to calculate the localization errors. The general structure of the automated methodology is presented in figure 1. The process can be summarized by a loop of four blocks: NSGA-II, Weighting, Source Reconstruction, and Performance indexes, where the outputs of NSGA-II are a set of best channel combinations and a set of all candidates.

**Figure 1:**
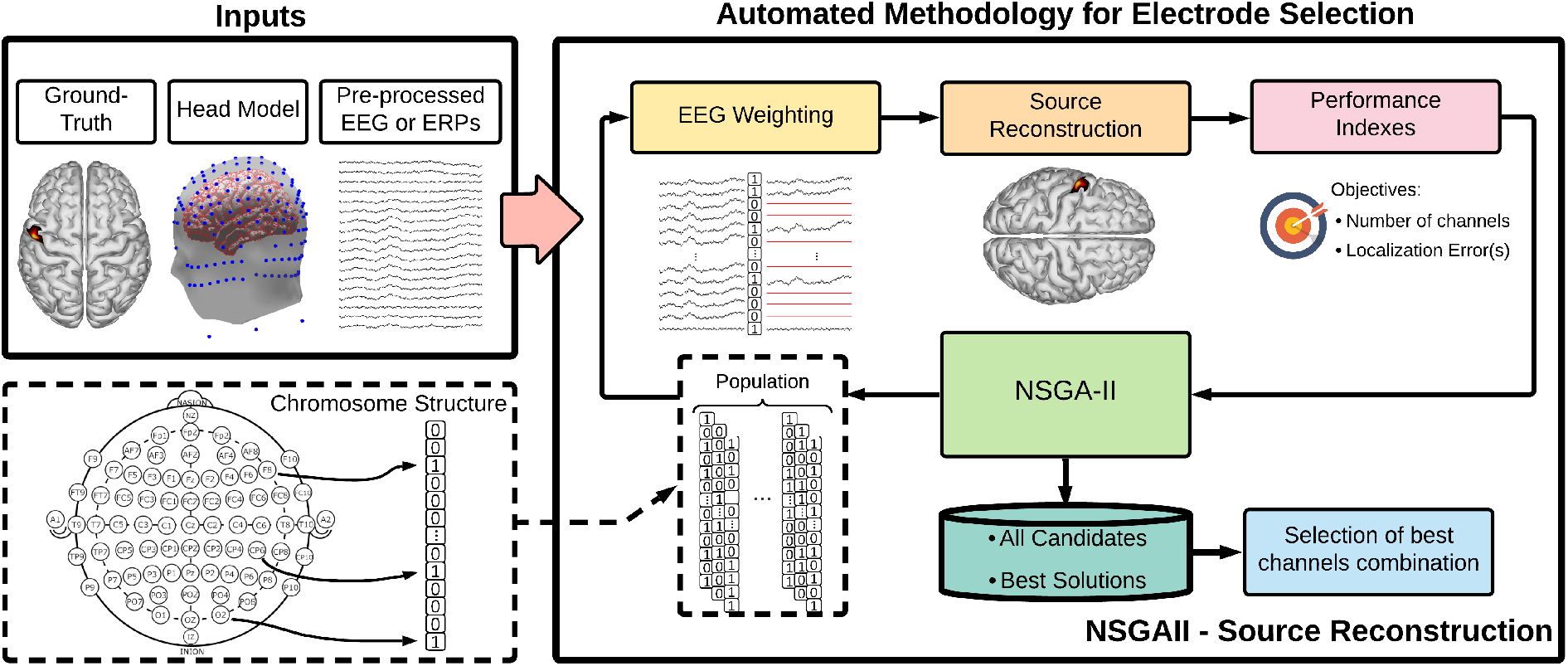
Flowchart of the proposed automated methodology for channel selection based on genetic algorithm multi-objective optimization.

An optimization problem consists of maximizing or minimizing a function by systematically choosing input values from a valid set and computing the value of the function, which can be limited to one or more restrictions, or it can be without any restriction. In an optimization problem, the model is feasible if it satisfies all the restrictions, and it is optimal if it also produces the best value for the objective function. In the case of a Multi-objective optimization problem (MOOP), it has two or more objective functions, which are to be either minimized or maximized. As in a single-objective optimization problem, a MOOP may contain a set of constraints, which any feasible solution must satisfy (Kalyanmoy Deb (2001)). In a multi-objective optimization problem, there is a set of solutions that is superior to the others in the search space when all the objectives are considered, but inferior to the other solutions for one or more objectives. Such solutions are known as Pareto-optimal solutions or *non-dominated solutions* and the rest as dominated solutions. The non-dominated sorting ranking selection method is used to emphasize good candidates and a niche method is used to maintain stable sub-populations of good points. NSGAs were created based on this concept (Srinivas and Deb (1994)).

The central block of NSGA-II consists of several stages: Population initialization, Fitness calculation, Crossover, Mutation, Survivor selection, and Termination criteria to return the best solutions. The population consists of a set of chromosomes, which are possible solutions to the problem, and each chromosome can have as many genes as variables in the problem. In this case, each gene represents an EEG channel, and the chromosome contains as many genes as the number of EEG channels in the registered EEG, represented by ***y***. A representation of the chromosome structure representing the electrode combination is presented at the bottom-left of figure 1. The NSGA-II initializes with a population of *P* chromosomes with random binary values. Then, in the weighting block, the genes values are used to weight the EEG by a dot multiplication between the EEG and the chromosome to compute the weighted EEG ***y_w_*** = ***y** · chromosome*. As a result of this mathematical operation, the channels with a gene value of one keep their value, while information of the channels with a gene value of zero is discarded.

The number of objectives *O* is defined as *O* = ***s*** + 1. Where, the algorithm considers a fixed objective to minimize **the number of EEG channels** used to localize single or multiple sources. The subsequent objectives are to minimize **the localization error** of the number of sources ***s*** desired to localize, with a separate objective per each source. NSGA uses a non-dominated sorting ranking selection method to emphasize good candidates and a niche method to maintain stable sub-populations of good points (Pareto-front), where a non-dominated solution is a solution that is not dominated by any other solution (Srinivas and Deb (1994)).

The weighted EEG ***y_w_*** is used in the source reconstruction block for calculating of the source activity and estimating the location of each source. In each weighted EEG ***y_w_*** the non-selected channels are converted to a temporal data series with zero value, therefore, the inverse solution is calculated using all the electrodes available in the head conduction model, but the information of contained in the non-used electrodes is discarded during source estimation. The use of all electrodes is preferred to avoid increasing the ill-posedness of the inverse problem when using a subset of the volume conduction matrix and to avoid using computational resources for re-calculating the volume conduction for each candidate combination.

In the performance indexes block, the number of channels extracted from each chromosome and the localization errors per each source are computed by comparing the estimated position with the ground-truth. The performance indexes return to the NSGA-II block that applies the non-dominated sorting, the half of the population with better performance is used to create the next generations using crossover and mutation procedures. The actual generation finishes when the new population is created and the process is repeated with the next generation until the termination criteria is reached. We used a maximum number of generations as criteria to stop the algorithm.

Finally, from the all set of chromosomes, the best combination per each number of channels is extracted to create a pseudo-Pareto front. In the case of the multiple sources, the pseudo-Pareto front is generated from the mean of the localization error between all the sources.

### 2.2. EEG Forward and Inverse Problems

The electrical activity recorded by the electrodes and its relation with the sources at cortical areas can be represented by the following equation, known as forward problem equation (Pascual-Marqui (1999)):

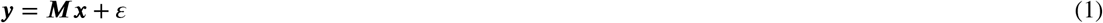

The forward problem allows to compute the potentials registered in the scalp, where the electrodes are placed, that were produced by current sources in the brain cortical areas. In the forward problem equation 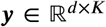 represents the signals recorded by *d* number of electrodes in *K* time samples; and 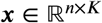 represents the time course activity of *n* sources at the cerebral cortex. The forward model 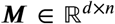 represents the relationship of the cortical source activity ***x*** with the electrodes measurements ***y***. This matrix is also known as lead field matrix or volume conductor model. In the EEG case, ***M*** represents how the electrical field propagates from the current sources in the brain to the scalp, where the voltages are registered by the electrodes.

The inverse problem is the estimation of the current sources in the brain from the electrical potential registered by the electrodes. Computing the sources from the electrode information is ill-posed due to the fact that the number of sources to estimate are much higher that the number of electrodes, an infinite combination of source activity could lead to the same recordings obtained by the electrodes. In addition, the problem is also ill-conditioned because the solutions are very sensitive to noise, and low perturbations on the channels can highly distort the source estimation (Grech et al. (2008)).

#### 2.2.1. Source Reconstruction Algorithms

One of the factors that influences the accuracy of source localization is the inverse algorithm used. For this reason and to consider this influence we have selected three widely used algorithms to solve the inverse problem: wMNE (Hämäläinen and Ilmoniemi (1984); Fuchs et al. (1994)), sLORETA (Pascual-Marqui (2002)), and MSP (Friston et al. (2008)).

wMNE is a variant of the original minimum norm estimation (MNE) (Hämäläinen and Ilmoniemi (1984, 1994); Dale and Sereno (1993)) based on Tikhonov regularization. wMNE involves a weighting matrix to compensate the deep sources distance to electrodes Iwaki and Ueno (1998); Pascual-Marqui (2002). sLORETA is a well-known method recognized by its zero localization error in absence of noise. This method is based on minimum norm, and improves the solution by standardizing the solution using the variance of the estimated activity Pascual-Marqui (2002). MSP is based on a Bayesian approach, and utilizes a set of priors over the cortical sheet to optimize hyper-parameters related to the covariance of the source activity Friston et al. (2008); López et al. (2014). This method is characterized to offer focalized (less blurry) estimation of the source activity.

#### 2.2.2. Source Reconstruction Error Measurements

The localization error is defined as the euclidean distance between the position in a 3D coordinated space of the ground-truth source *P_x_* and the estimated source position 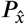 by:

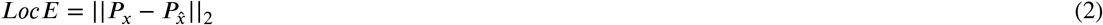

This error was used in the proposed methodology as objective to minimize. Relative error and Pearson Correlation Coefficient were used as additional error measurements to compare the time courses between reconstructions done with different number of electrodes. Consider *x*_1*i*_ and *x*_2*j*_ as the time courses of two reconstructions in the vertices *i* and *j* were the source with highest amplitudes were found, being *x*_1*i*_ the time course of the reconstruction computed with the highest number of electrodes. These metrics can be computed by using the following equations:

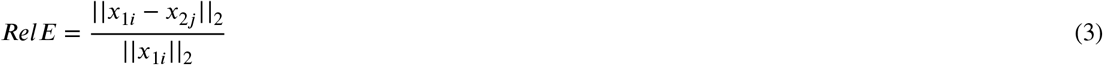

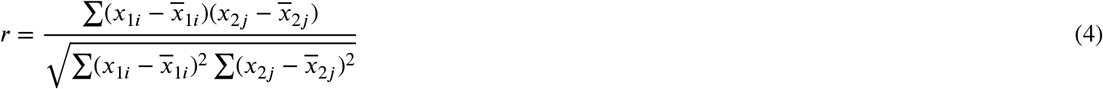

where 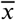 represents the mean of the values of *x*.

### 2.3. Evaluation Framework

We evaluated the proposed methodology over synthetic and real EEG signals. In the section below, we present the synthetic EEG dataset with multiple sources activity and its simulation framework that can be used for evaluating inversion algorithms. We have tested the automated methodology for channel selection over this dataset, under single and multiple sources cases. Furthermore, we have applied the methodology on real EEG signals, by using the dataset “Localize-MI” presented in Mikulan et al. (2020), where HD-EEG was recorded during intracerebral stimulation, and the ground-truth activity location is therefore available. Multiple tests of the methodology have been perform along this document and they are presented and described in section 2.3.3

#### 2.3.1. Synthetic EEG Dataset

The synthetic EEG dataset consists of 150 trials of multiple source activity. We used the forward equation 1 to simulate the EEG data. Per each trial the sources were distributed over several regions of the brain, the time courses of the sources were generated using a Gaussian windowed sinusoidal activity as in Soler et al. (2020b), defined by the following equation:

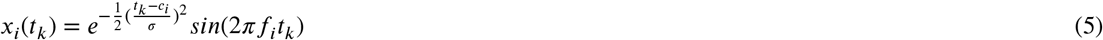

where *i* represents the source number, *c_i_* the time center of the source activity represented in seconds, *σ_i_* determines the shape of the Gaussian window by adjusting its width, and *f_i_* is the frequency of the sinusoidal activity. The first source *s*_1_ was simulated in the occipital lobe, the activity was centered at *c*_1_ = 0.5*s*, and the frequency activity set at *f*_1_ = 19*Hz* in the range of beta rhythm. The second source *s*_2_ was located in the sensory-motor cortex, with a time center of *c*_2_ = 1*s*, and a frequency of *f*_2_ = 10*Hz* in the range of the mu rhythm. The third source *s*_3_ was simulated with a frequency of *f*_3_ = 7*Hz* in the range of theta rhythm, it was centered in time at *c*_3_ = 1.5*s*, and located in the frontal lobe. Additional three sources, *s*_4_, *s*_5_ and *s*_6_ were generated with similar parameters and centered at time *c*_4_ = 2*s*, *c*_4_= 2.5*s*, and *c*_6_ = 3*s* but they were not used for testing in this study, therefore we refer as all sources for only the first three sources of the dataset. An example of the time course of the simulated sources and their location is shown in figure 2. For all the sources the parameter window width was set at *σ* = 0.12, this value combined with the center of each source allowed to have a temporal mixing between sources, this temporal mixing is represented with the light-gray areas in figure 2. Notice that source *s*_1_ only has temporal mixing in the transition to the next source, but *s*_2_ and *s*_3_ have temporal mixing in both sections of the waveform, where around 40% of the source activity is overlapped.

**Figure 2:**
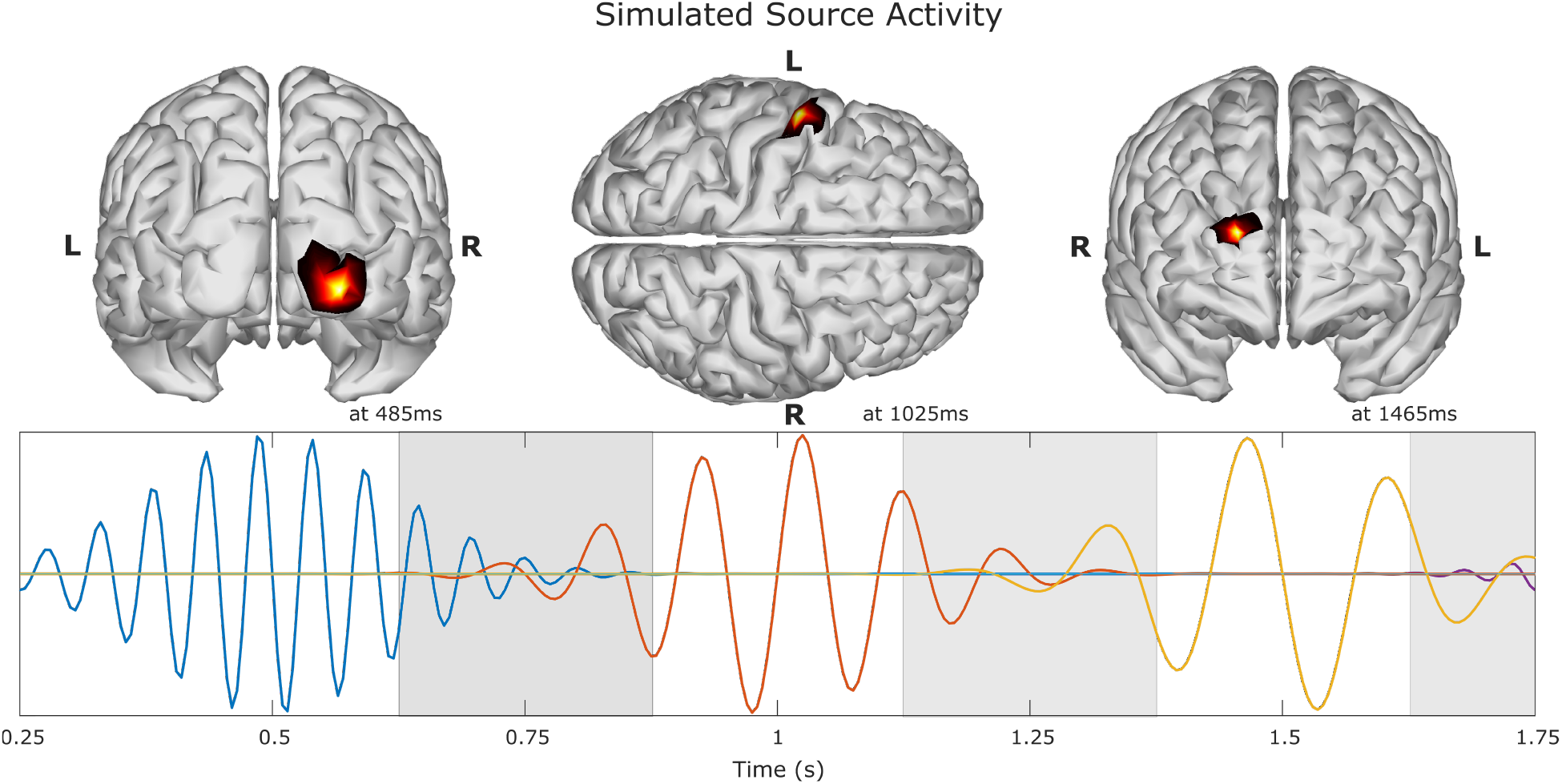
Example of simulated source activity. Source *s*_1_ time course is depicted in blue color, source *s*_2_ in orange, source *s*_3_ in yellow, and the remaining activity from *s*_4_ in purple. The locations of the first three sources are depicted on top, where the time was chosen where the maximum amplitude value for each source took place. The light-gray areas depict the time sections where there are temporal mixing between sources.

The sources were distributed over the brain in three main areas, occipital lobe, sensory-motor cortex, and frontal lobe. Per each area, a set of twelve positions (six per hemisphere) was predefined, and during the simulation procedure the position was randomly selected from the pre-defined sets. This allowed to create trials with different combinations of sources. Figure 3 shows the locations of the pre-defined sources (represented by the blue circles), and the number of times they were selected in the dataset (represented by the diameter of the circles).

**Figure 3:**
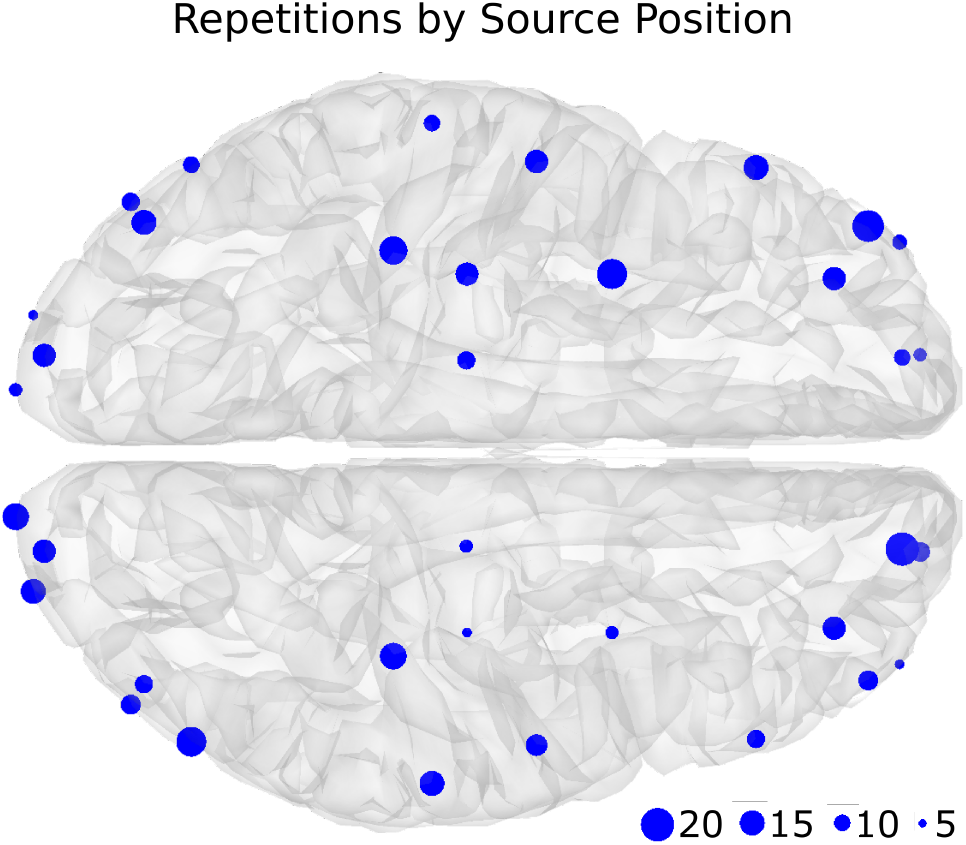
Repetition by source position. Blue circles represent where the simulated activity was placed, and their diameter represents the number of times the position was selected in the 150 EEG trials. It can be seen that each area: occipital lobe, sensory-motor cortex, and frontal lobe has six locations by hemisphere.

The lead-field matrix for solving the forward problem was taken from a FEM forward model called “New York head model” (Huang et al. (2016)). The model is available in https://www.parralab.org/nyhead/, it was computed modelling the scalp, skull, air cavities, CSF, gray matter, and white matter. In FEM models, the influence of including the CSF and the white matter during modelling have been found significant for solving the inverse problem (Vorwerk et al. (2014)). The New York head model was proposed as a standard model to be used in EEG studies, and it is based on a non-linear average of the MRI of 152 adult human brains. The forward model includes a high-resolution lead field matrix relating 75000 sources with 231 channels, it also includes additional versions with a reduced number of sources of 10000, 5000 and 2000 sources. The model considers 231 electrodes, from which 161 are located in the scalp (based on the 10-10 and 10-5 systems), 2 in the left/right pre auricular point (LPA/RPA), 4 in the neck, and 64 distributed around the face and in the back of the head below Iz channel. Figure 4 presents a lateral and top view of the channels and sources locations using the model with 10000 sources.

**Figure 4:**
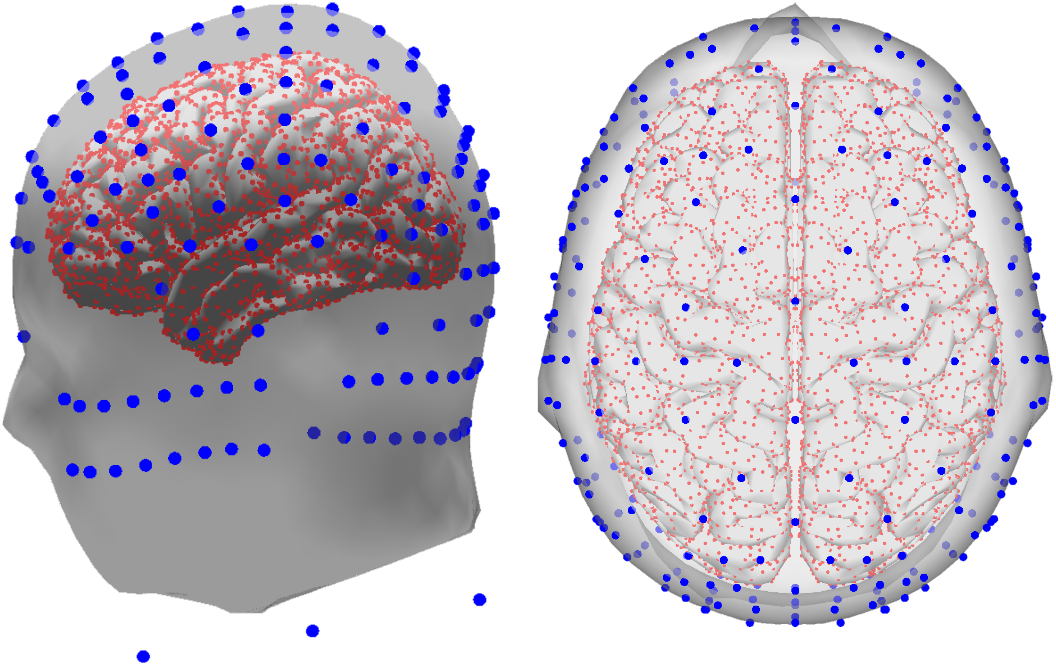
New York head model with 10000 sources and 231 channels (Huang et al. (2016)). Lateral view (left), and top view (right). The blue circles represent the location of the channels over the scalp, face, and neck, while the red circles, the location of the distributed sources on the brain’s cortical areas.

The model of 10000 sources was used for forward computations to generate the EEG signals 2.3.1, and the model of 5000 sources to calculate the inverse solutions. It allows to avoid the effects of the so-called inverse crime (Colton and Kress (2019)). The use of the New York head model Huang et al. (2016) in our experiments is based on several reasons: one reason is that the model was computed using FEM including six tissues, in FEM models the influence of including the CSF and the white matter during modelling have been found significant for solving the inverse problem (Vorwerk et al. (2014)), using a more detailed model can yield to more accurate source reconstruction results. Another reason was the number of electrodes available in the model, it considers 231 channels located as per the standard 10-20 and 10-10, and the extended system 10-5. It is well known that more channels lead to a better reconstruction Sohrabpour et al. (2015); Song et al. (2015), therefore for a better comparison between the proposed methodology (that identifies an optimized subset of channels) and the full set of electrodes, using the maximum posible number of electrodes is desirable.

After calculating the EEG using equation 1, the signals of the channels were corrupted with noise with a signal-to-noise ratio (SNR) of 0*dB* (signal and noise with the same power). Each trial has a duration of 3.5*s*, with a sampling frequency of 200*Hz*. The dataset with the 150 trials and ground-truth information is available in https://github.com/anfesogu/Ground-Truth-EEG-Dataset to facilitate replication of the results in this paper. It can also be used in future studies where a ground-truth for source localization is required.

#### 2.3.2. Real EEG Dataset

We used the dataset “Localize-MI” by Mikulan et al. (2020) to test the methodology over real EEG signals. The dataset consists of HD-EEG recordings and anonymized MRI of seven participants that were stimulated with single-pulse biphasic currents by implanted electrodes, the stimulation position is known and the dataset serves as ground-truth for evaluating inverse solutions. The EEG signals were recorded with 256 geodesic electrode system (Geodesic Sensor Net; HydroCel CleanLeads) where the electrodes position were digitized to allow co-registration with the MRI. Referring to the original dataset publication: “All of the participants provided their Informed Consent before participating, the study was approved by the local Ethical Committee (protocol number: 463-092018, Niguarda Hospital, Milan, Italy) and it was carried out in accordance with the Declaration of Helsinki.” from Mikulan et al. (2020)

The dataset contains a set of pre-processed epochs from −300 to 50 ms around the stimulation artifact. They were filtered with a high-pass filter at 0.1 Hz, and notch filter at 50, 100, 150, and 200 Hz (only subjects 5 and 7). Baseline correction was applied between −300 to −50. Finally the epochs were averaged and cropped 2*ms* around the stimulation artifact. In total, the dataset consist of 61 sessions, all of them were further used for testing the methodology. We refer to the original dataset publication for more detailed information (Mikulan et al. (2020)).

A head model of each participant was created to be used during source reconstruction. The models were created by processing the individual MRI using Freesurfer (Martinos Imaging Centre, Boston, MA, Fischl (2012)) and MNE-python (Gramfort et al. (2014)). The number of sources was defined as 4098 per hemisphere, and the lead field matrix was computed using BEM, considering the conductivities of scalp, skull, and brain as 0.3, 0.006 and 0.3*S/m* respectively (MNE-python default values).

#### 2.3.3. Test Structure

In this study we proposed multiple test to investigate to what extend the quality of the source reconstruction of HD-EEG, measured in terms of source location error, can be maintained while using a set of reduced number of channels that were selected by the proposed methodology. The first test was based on source reconstruction for one source, we refer to it as “single-source test”. This test was performed by setting two minimizing objectives for the optimization: the localization error of the single source and the number of channels. We considered the first source *s*_1_ of the synthetic dataset, the epoch was set between 250 to 750 ms, including a section of overlapping with the next source *s*_2_.

A second test was done considering the first three sources of the synthetic dataset *s*_1_, *s*_2_, and *s*_3_, we refer to it as “multiple-source test”. In this test, the epoch definition was set between 250 to 1750 ms, and four minimization objectives were set for the optimization algorithm: the individual localization error of each source and the number of channels. To evaluate the approach over different number of electrodes, the multiple-source test was performed over the full set of 231 electrodes, we refer at it as “multiple-source test 231e” (161 electrodes at scalp, of those 73 are located in positions of the standard 10-10 system and 88 located in positions of the standard 10-5 system, the other electrodes are placed in the neck and face areas (Figure 4)). A second test was performed constraining the search space for the optimization algorithm to 128 scalp electrodes, of which 64 channels were selected from the standard 10-10 and 64 channels from the standard 10-5, we refer at it as “multiple-source test 128e”. Finally a third test was performed constraining the search space of the NSGA-II to 60 electrodes, all of them located in standard 10-10 positions, this test is referred as “multiple-source test 60e”. In figure 5. is shown the 161 positions of the scalp electrodes of the New York head model, and the subsets configuration for 128 and 60 electrodes.

**Figure 5:**
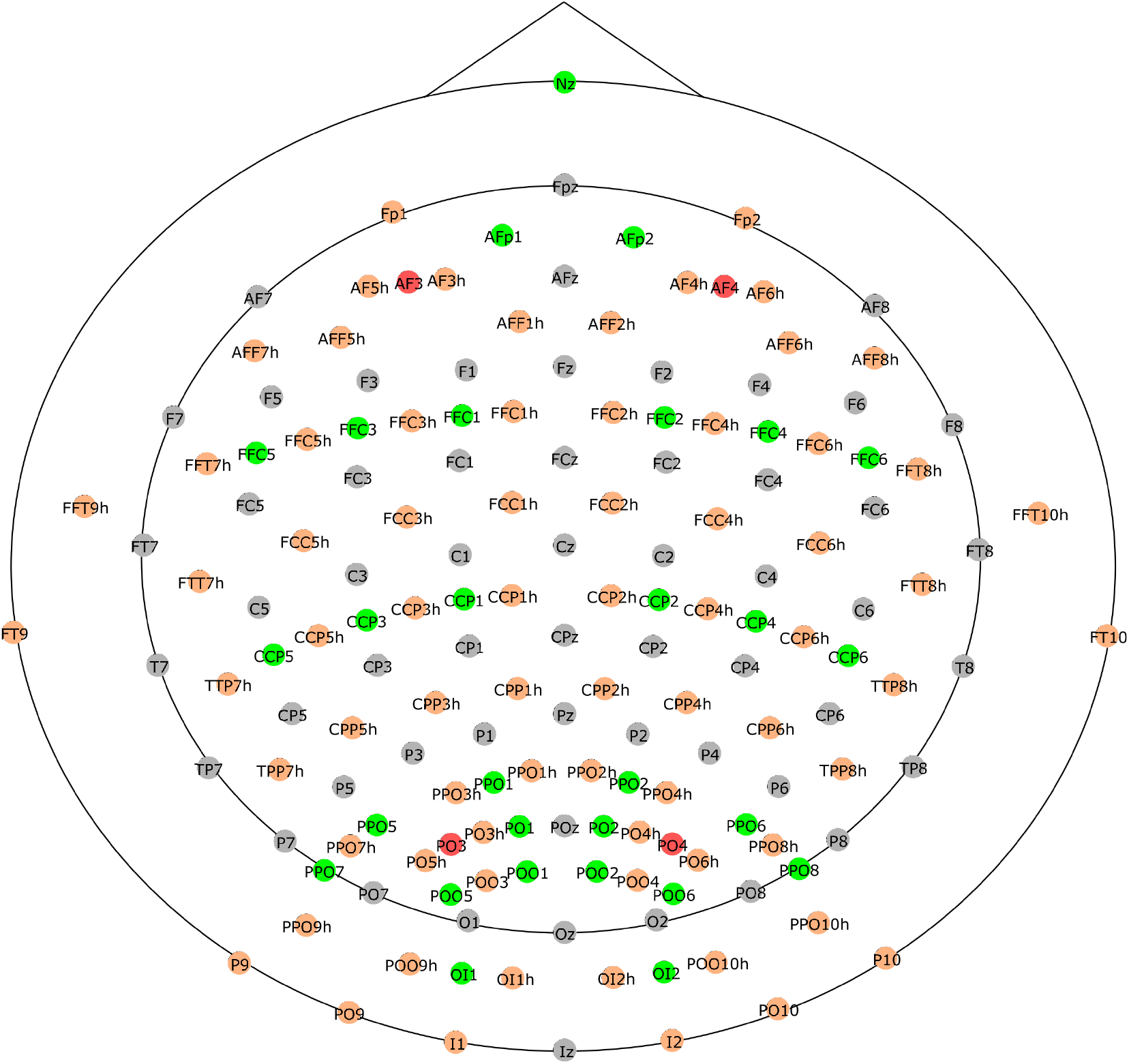
Location and names of the 161 scalp electrodes included in the New York head model. Red and gray circles represent the subset of 60 electrodes, and orange and gray circles represent the subset of 128.

The third test was done using the Localize-MI dataset. During each session the stimulation took place in the same electrode position, therefore, we treated the data as single source case, and we set two minimizing objectives for the optimization algorithm: the localization error of the artifact source and the number of channels. We refer to this test as “Localize-MI test”.

The procedure during all tests started by defining the electrodes to be considered, only in the multiple-source tests the electrode number varied according to the test (when constraining to 128 or 60 electrode positions); in the single-source test all the electrodes were considered. In the case of the Localize-MI test, the electrodes that were marked as bad were removed when calculating the forward model, no attempt to clean or interpolate channels was carried out. The number of electrodes to use defines the length of the chromosome for the optimization. After setting the number of electrodes, the number of objectives is defined, and the EEG of each trial (in the case of the synthetic data) or the ERP of the session (in the case of the Localize-MI dataset) is transferred to the algorithm. In the next step, the population size and the maximum number of generations are set, their values were determined experimentally and set as 100 and 400 respectively. Finally, the algorithm for source reconstruction is defined, and the algorithm can start processing data by combining the NSGA-II with sLORETA, wMNE and MSP in separated runs respectively.

Each trial or ERP is processed by the algorithm at least three times, each time with a different source reconstruction algorithm. During a run, the algorithm evaluates 40000 combinations of electrodes while trying to minimize the objectives. When the run finalizes, a output file with the combinations and performance indexes of all generations and chromosomes is generated, from it, the performance of the best combination per each number of channels is extracted to create the pseudo-Pareto front. Finally an accuracy index was computed for comparing the reconstructions from the optimized set of channels versus all the channels available, the accuracy index represents the percentage of trials or ERPs that obtained equal or lower localization error with a given number of electrodes when comparing the accuracy when using all the electrodes for the same trial or ERP.

The proposed methodology was implemented and executed in Matlab (The MathWorks, Inc.) version 2016. The source reconstruction by wMNE and sLORETA were implemented as custom functions in Matlab. To run the MSP method, we used the Statistical Parametric Mapping (SPM; Wellcome Centre for Human Neuroimaging, London, UK) software for Matlab. The NSGA-II solver by Song (2011) was adapted to include source reconstruction. The tests were carried out using the IDUN computing cluster (Själander et al. (2019)) of Norwegian University of Science and Technology NTNU.

## 3. Results

### 3.1. Single-source test

The pseudo-Pareto front from the single-source test and localization error with all the electrodes are presented in figure 6. The pseudo-Pareto front exhibits a typical convex behavior, the accuracy obtained with the optimized combinations of channels presents a stable value for the localization error when using combinations with five or more channels, independently of the source reconstruction method applied. It can be seen an inflection point in the pseudo-Pareto front when reducing from five to four channels, it is also evident when looking at the values presented in table 1, where the results from the optimization with two to sixteen channels and the results with all the electrodes are summarized. The accuracy index of the optimized combinations of channels was extracted by quantifying the percentage of trials with a given number of channels that obtained equal or lower localization error than when using all the electrodes for the same trial. The accuracy index for the single source test exhibits a relatively stable behavior from 16 to 5 channels, but it has a decrease of more than two perceptual points when reducing from five to four channels. Its value continues decreasing substantially while reducing the channels, coinciding with the increasing of the localization error and the standard deviation.

**Figure 6:**
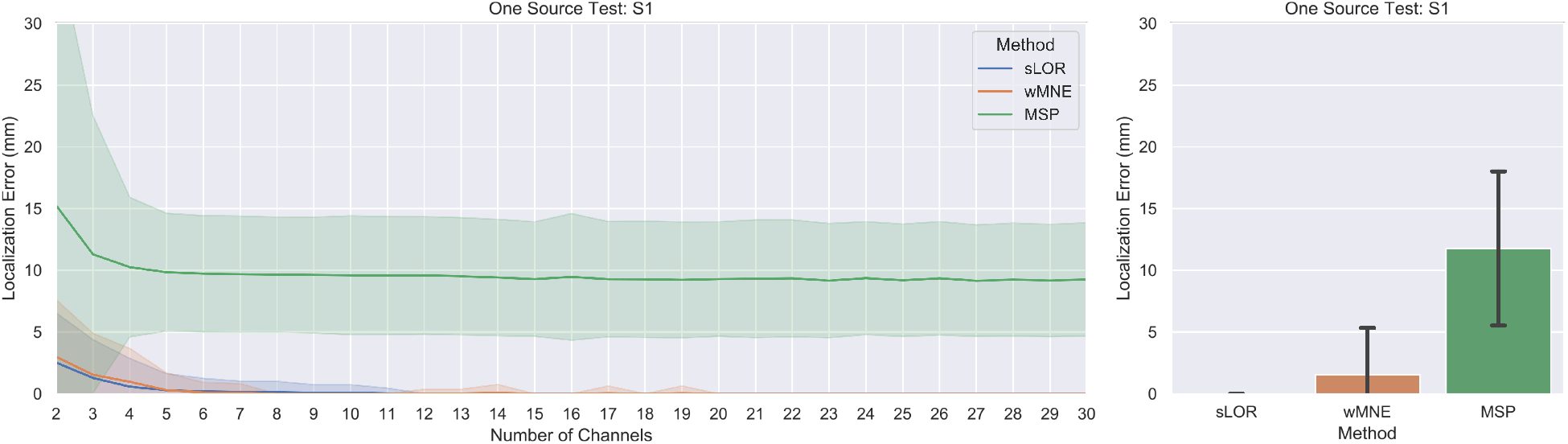
Results for the single-source test. The pseudo-Pareto fronts relating the number of channels and the localization error by each source reconstruction method are presented at left. The lines represent the mean localization error across the 150 trials and their respective colored bands represent their standard deviation. The mean localization errors across the trials and the standard deviation obtained by using all the electrodes available (231) by each method are presented at right.

**Table 1.**
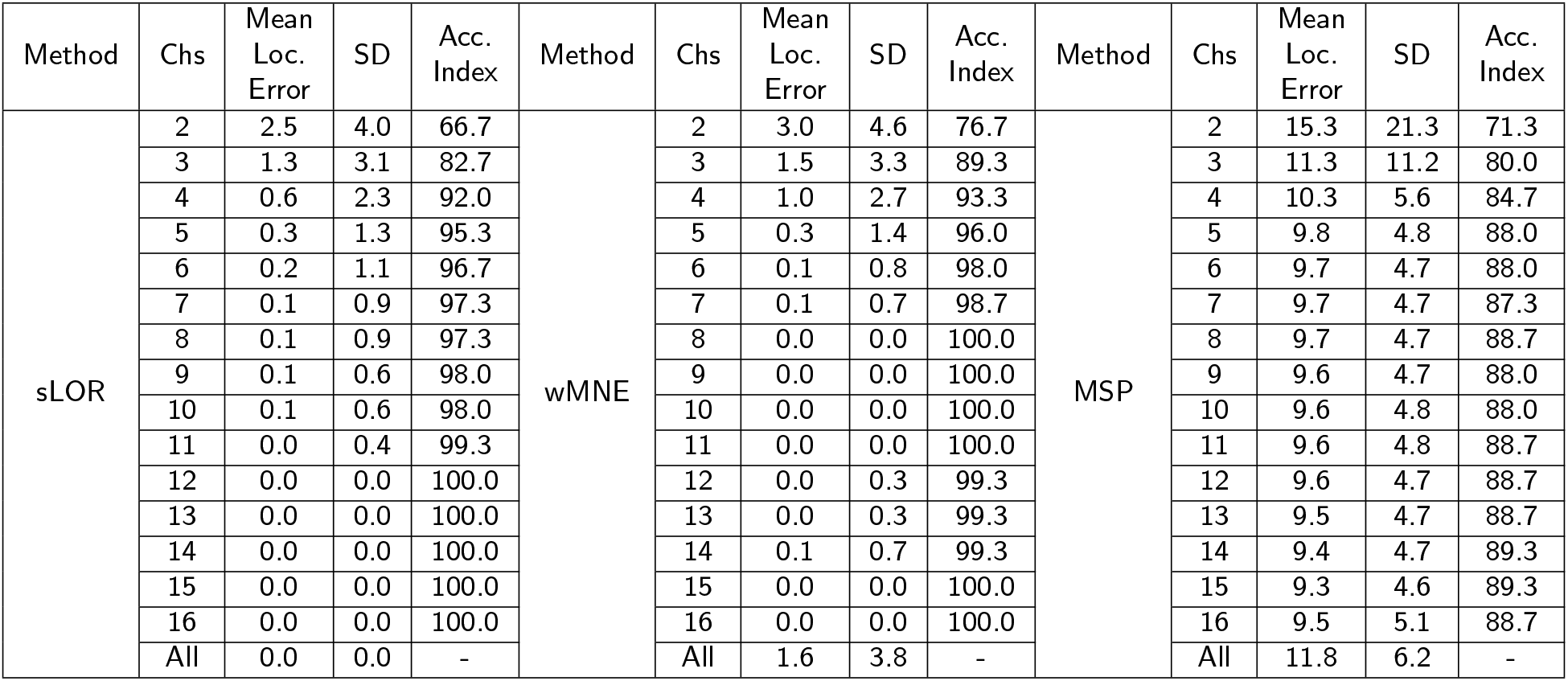
Summary of the localization error obtained with the optimized combinations of channels, from 2 to 16, and with the full set of 231 electrodes. The accuracy index represents the percentage of trials with the given number of channels that obtained equal or lower localization error than when using all the electrodes for the same trial.

When analyzing the results presented in figure 6 and table 1 in a more detailed level, it can be seen that with a combination of four or more channels, more than 92% of accuracy index was obtained for sLORETA and wMNE, indicating that at least 138 over 150 trials obtained equal or lower accuracy than when using all the electrodes. In the case of MSP it presents a lower accuracy index than the other methods, with a stable value around 88%, and when considering combinations with four or more channels at least 127 of 150 trials were equal or better than the reconstructions with all electrodes.

### 3.2. Multiple-source test

The pseudo-Pareto Front for the multiple-source test with the different electrode configurations are shown in figure 7. The multiple test had four optimization objectives. To facilitate the visualization of the results, the pseudo-Pareto front was obtained from the mean localization error across the 150 trials, where the localization error is the average of the individual localization error for each one of the three sources. Table 2 presents the localization error, standard deviation and accuracy index for the multiple-source tests for a selected number of channels and with all electrodes.

**Figure 7:**
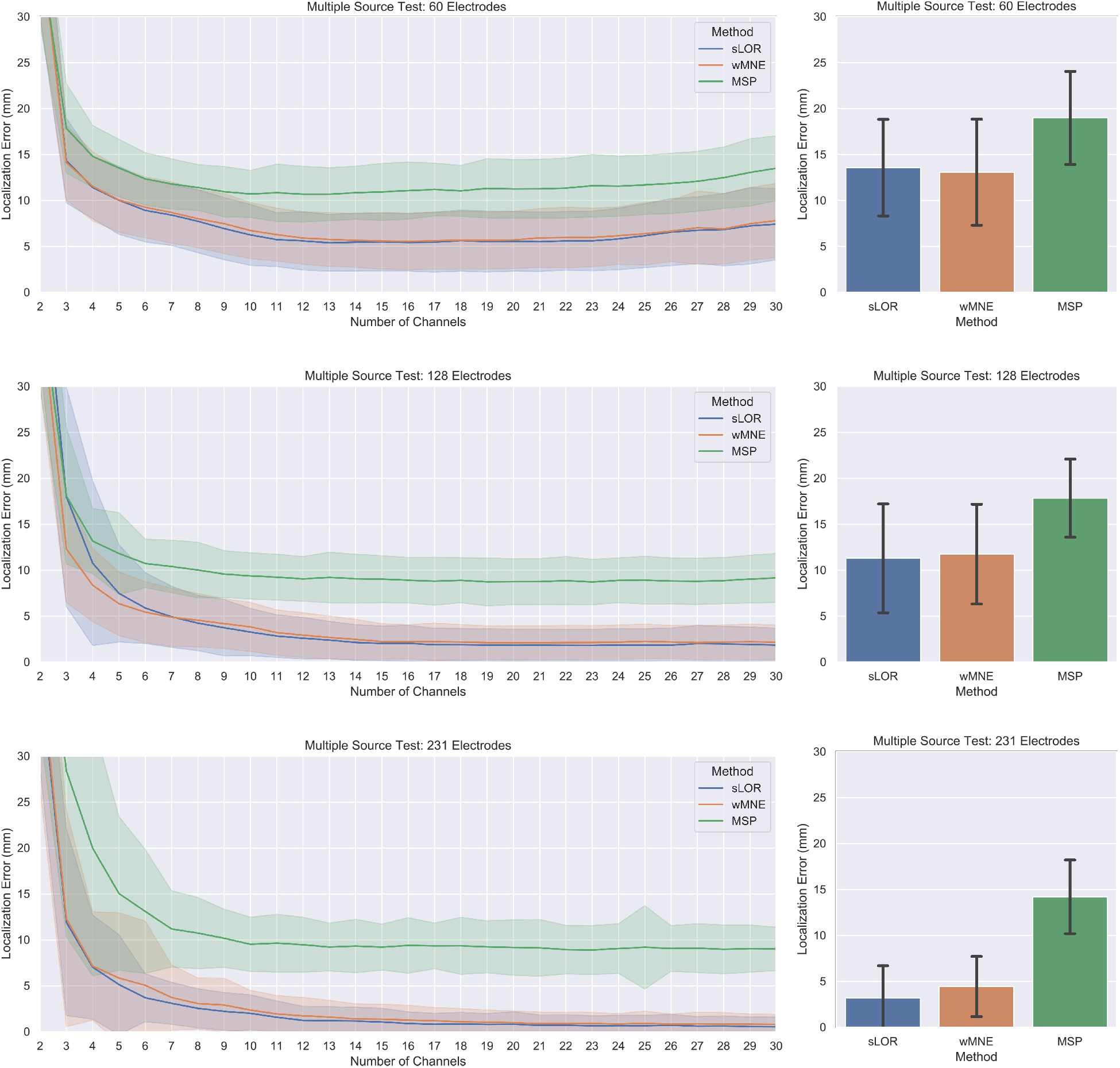
The pseudo-Pareto fronts for the multiple-source test with the different constrained number of channels are shown at Left, they relate the number of channels vs the average between the three sources localization error. The lines are representing the mean of the localization error* of the three sources across the 150 trials, and the colored bands the standard deviation. The mean localization error* across the trials and the standard deviation obtained by using the subsets of 60 and 128 electrodes, and the full set of 231 electrodes for the three sources are presented at right. * The mean localization error is computed using the average error between the three sources and then the average of this value across the 150 trials.

**Table 2.** Multiple-sources test results.

| Method | Chs | Mean Loc. Error | SD | Index (%) | Method | Chs | Mean Loc. Error | SD | Index (%) | Method | Chs | Mean Loc. Error | SD | Index (%) |
| --- | --- | --- | --- | --- | --- | --- | --- | --- | --- | --- | --- | --- | --- | --- |
| sLOR Test 60e | 4 | 11.40 | 3.38 | 65.33 | wMNE Test 60e | 4 | 11.53 | 3.75 | 58.00 | MSP Test 60e | 4 | 14.79 | 3.36 | 74.00 |
|  | 6 | 8.91 | 3.43 | 84.67 |  | 6 | 9.25 | 3.30 | 74.67 |  | 6 | 12.34 | 2.85 | 90.67 |
|  | 8 | 7.73 | 3.42 | 91.33 |  | 8 | 8.01 | 3.15 | 84.67 |  | 8 | 11.40 | 2.49 | 97.33 |
|  | 10 | 6.25 | 3.31 | 98.00 |  | 10 | 6.72 | 3.05 | 92.00 |  | 10 | 10.70 | 2.56 | 98.67 |
|  | 12 | 5.59 | 3.16 | 98.67 |  | 12 | 5.90 | 2.85 | 96.00 |  | 12 | 10.68 | 3.02 | 98.00 |
|  | 14 | 5.45 | 3.15 | 98.67 |  | 14 | 5.64 | 2.91 | 96.67 |  | 14 | 10.83 | 2.88 | 98.00 |
|  | 16 | 5.44 | 3.12 | 99.33 |  | 16 | 5.55 | 3.08 | 98.67 |  | 16 | 11.07 | 3.10 | 97.33 |
|  | All | 13.56 | 5.25 | - |  | All | 13.06 | 5.77 | - |  | All | 18.98 | 5.07 | - |
| sLOR Test 128e | 4 | 10.75 | 8.92 | 55.33 | wMNE Test 128e | 4 | 8.35 | 3.96 | 69.33 | MSP Test 128e | 4 | 13.16 | 3.54 | 74.67 |
|  | 6 | 5.87 | 3.84 | 79.33 |  | 6 | 5.43 | 3.30 | 87.33 |  | 6 | 10.72 | 2.64 | 94.00 |
|  | 8 | 4.23 | 3.00 | 87.33 |  | 8 | 4.54 | 2.90 | 90.00 |  | 8 | 10.00 | 3.03 | 96.67 |
|  | 10 | 3.26 | 2.56 | 94.00 |  | 10 | 3.82 | 2.67 | 96.00 |  | 10 | 9.37 | 2.51 | 97.33 |
|  | 12 | 2.58 | 2.25 | 95.33 |  | 12 | 2.92 | 2.43 | 97.33 |  | 12 | 9.04 | 2.43 | 96.67 |
|  | 14 | 2.13 | 1.94 | 98.00 |  | 14 | 2.46 | 2.18 | 99.33 |  | 14 | 9.06 | 2.66 | 98.00 |
|  | 16 | 2.03 | 1.96 | 99.33 |  | 16 | 2.20 | 1.87 | 100 |  | 16 | 8.90 | 2.47 | 98.00 |
|  | All | 11.28 | 5.93 | - |  | All | 11.74 | 5.43 | - |  | All | 17.82 | 4.25 | - |
| sLOR Test 231e | 4 | 7.08 | 5.75 | 20.00 | wMNE Test 231e | 4 | 7.18 | 5.87 | 24.00 | MSP Test 231e | 4 | 20.15 | 13.81 | 29.33 |
|  | 6 | 3.69 | 2.59 | 44.00 |  | 6 | 5.06 | 6.99 | 47.33 |  | 6 | 13.15 | 6.76 | 56.00 |
|  | 8 | 2.54 | 2.13 | 58.00 |  | 8 | 3.06 | 2.78 | 60.67 |  | 8 | 10.74 | 3.87 | 74.00 |
|  | 10 | 2.00 | 2.00 | 71.33 |  | 10 | 2.35 | 2.14 | 66.67 |  | 10 | 9.53 | 2.99 | 82.00 |
|  | 12 | 1.23 | 1.53 | 88.00 |  | 12 | 1.72 | 2.00 | 76.00 |  | 12 | 9.47 | 2.97 | 81.33 |
|  | 14 | 1.16 | 1.49 | 90.00 |  | 14 | 1.38 | 1.64 | 82.00 |  | 14 | 9.32 | 2.90 | 84.00 |
|  | 16 | 0.90 | 1.26 | 96.67 |  | 16 | 1.26 | 1.60 | 82.67 |  | 16 | 9.39 | 3.02 | 86.00 |
|  | All | 3.17 | 3.52 | - |  | All | 4.43 | 3.28 | - |  | All | 14.19 | 4.01 | - |

The results for all electrodes in the three test show a tendency of increasing the localization error when using a lower number of electrodes (Figure 7 right column). When localizing the sources, the mean between 60 and 128 channels presents a similar value in each method, being the solution with 60 channels less accurate. In contrast, the difference is notorious when increasing from 128 to 231 channels, and it is more evident for sLORETA and wMNE methods. In the pseudo-Pareto’s fronts of the proposed methodology, the same tendency as with all electrodes can be observed. In them, the localization error was more accurate when using the full set of 231 channels, and when restricting the search space to 128 and 60 channels, the localization error increased. However, when comparing the solutions for the test with 128 and 231 channels, there is not considerable difference between the solutions, but the obtained localization error values were lower for sLORETA and wMNE in the multiple-source test with 231 electrodes.

When analyzing the values of table 2, it can be seen that for the multiple-source tests of 60e and 128e, the proposed methodology obtained a lower mean localization error when considering optimized combinations with four or more electrodes than with the 60e and 128e subsets. In the case of the multiple-source test 231e, the proposed methodology obtained lower errors than with all the 231 channels, when using optimized combinations of eight or more channels. Regarding the accuracy index, in the multiple-source test 231e, for the optimized combinations with 16 electrodes, in 145 of 150 trials the proposed methodology obtained equal or lower accuracy than the full set of electrodes. The accuracy index reduces as the number of channels decreases, at the point of eight electrodes the accuracy index presents a value of 58% (87 of 150 trials), however, despite the relative low accuracy index, the mean across the trials was lower than with the full set.

An example of the position of the true and estimated sources locations, and the selected channels are show in figure 8, it shows the optimized combinations with four and eight electrodes for one trial of the dataset, and the resulted combinations were obtained from the multiple-source test 231e results for each source reconstruction method. In the combinations of four channels, it can be observed that the channels selected are located close to the true source location, where the mean localization errors were 5.3, 4.7, and 14, 5 mm, for sLORETA, wMNE, and MSP, respectively. In addition, when observing the combination of eight channels, the selected electrodes were located not only close to the source location, also in intermediate positions between the sources that were relatively close. In contrast, for the sources that were more separated, there was not selected any channel between them. The mean localization errors for the eight channels combinations were 2.4, 4.2, and 4, 5 mm, for sLORETA, wMNE, and MSP, respectively. This suggest that the proposed methodology selected not only the channels that contained the most information from a single source, but also the channels that contained shared information from multiple sources.

**Figure 8:**
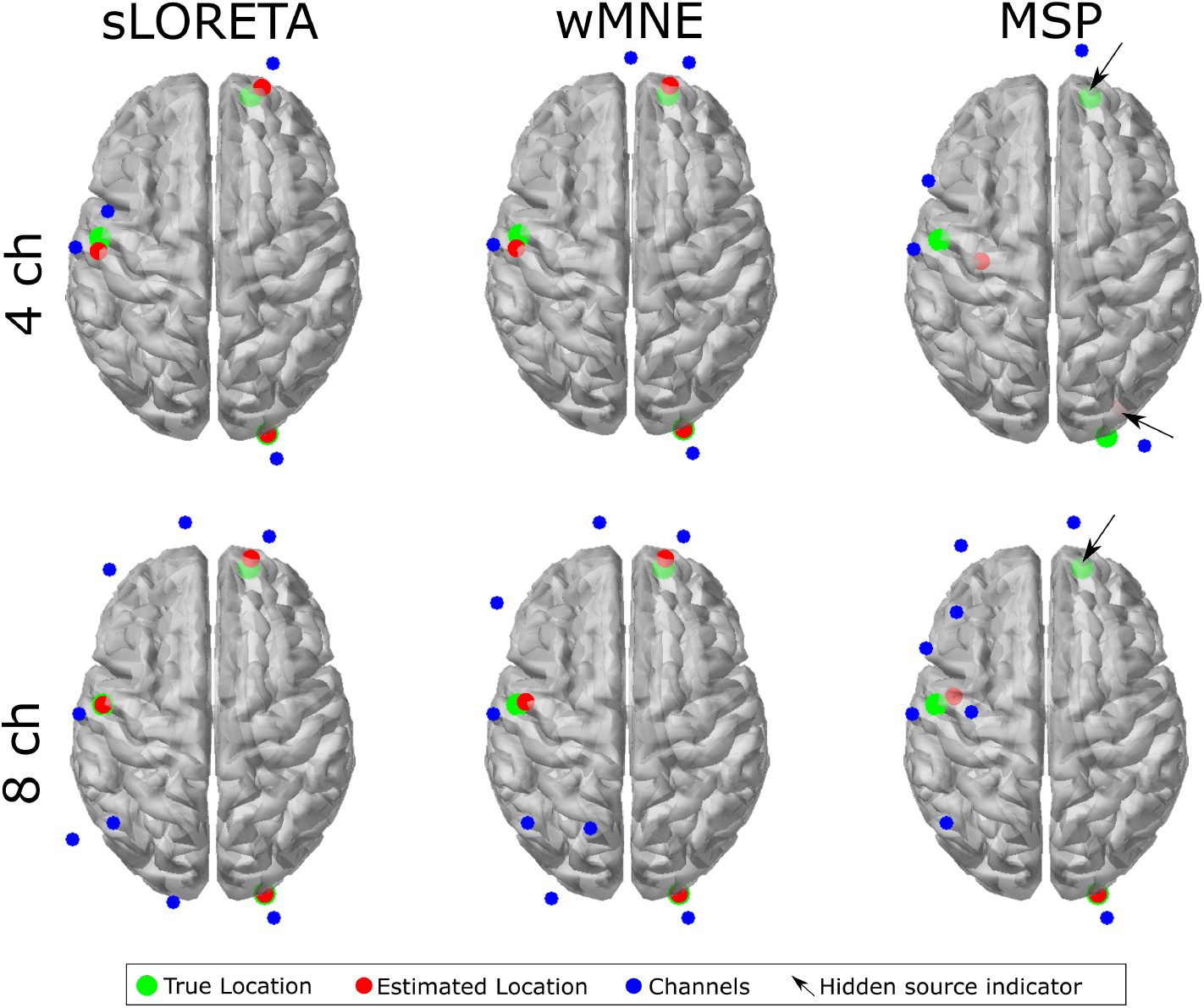
Optimized combinations and source localization for one trial during the multiple-source test 231e. The mean localization errors with four channels were 5.3, 4.7, and 14, 5mm, and with eight channels were 2.4, 4.2, and 4, 5mm, for sLORETA, wMNE, and MSP, respectively. The hidden source indicator is intended to point where there is a source that can not be seen from the top view.

Regarding the effects of the estimated waveform for each source, the standardized time courses of the sources estimated with 231 channels, and with the optimized sets of four and eight channels for the one trial are presented in figure 9. The sources waveform are highly similar independently of the number of channels, however, the reconstructions were noisier when using the optimized sets than with the full set of electrodes. For the same trial, the time courses of the pseudo-Pareto channel combinations were compared against the time course using the 231 channels, the relative error and Pearson correlation coefficient were calculated for each source, each number of channels from 3 to 30, and for each reconstruction algorithm, its values are presented in figure 10. It was found that the source time courses estimated with the optimized combinations and the full set had more than a 97% of correlation when considering five or more channels, obtaining a relative error lower than 0.25. When considering optimized combinations with more electrodes, the difference became smaller, the level of correlation with 16 channels or more is around 99%, independently of the source reconstruction method.

**Figure 9:**
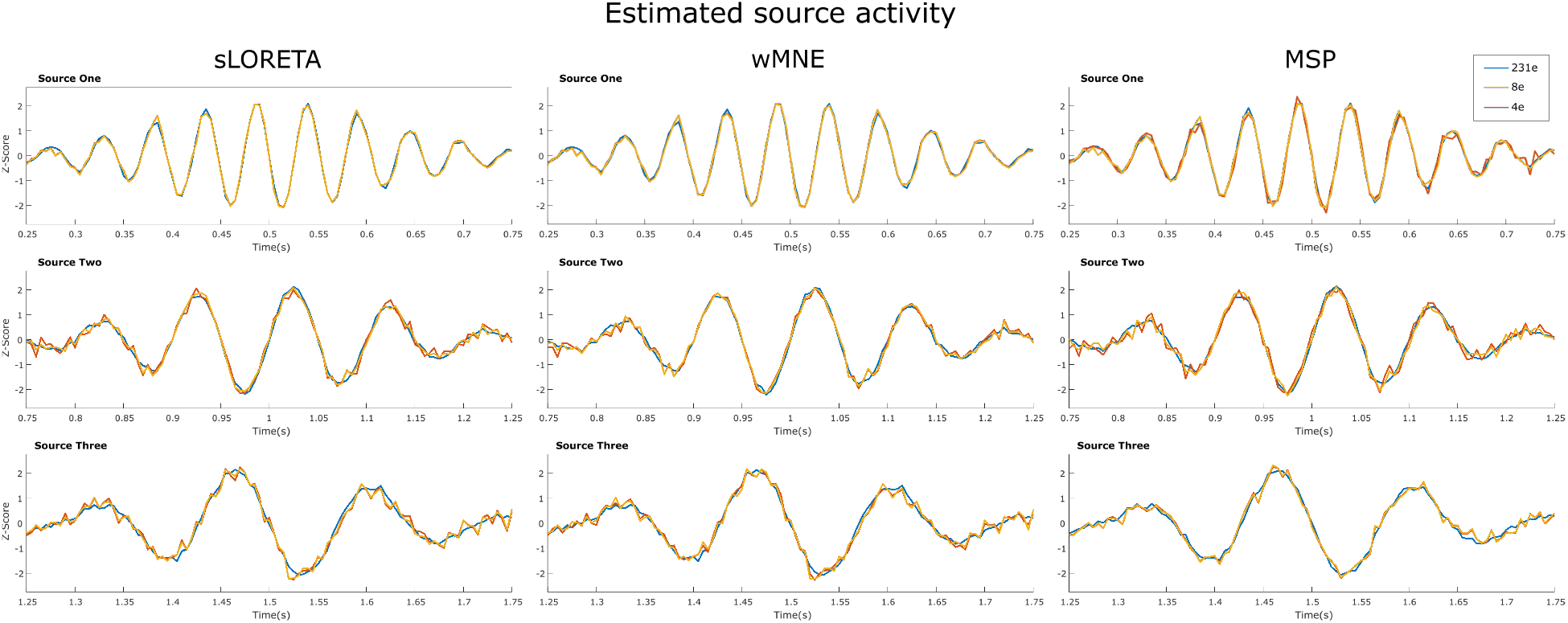
Example of the estimated time courses of the three sources with the full set of 231 channels, and with the optimized combinations with four and eight channels for one trial.

**Figure 10:**
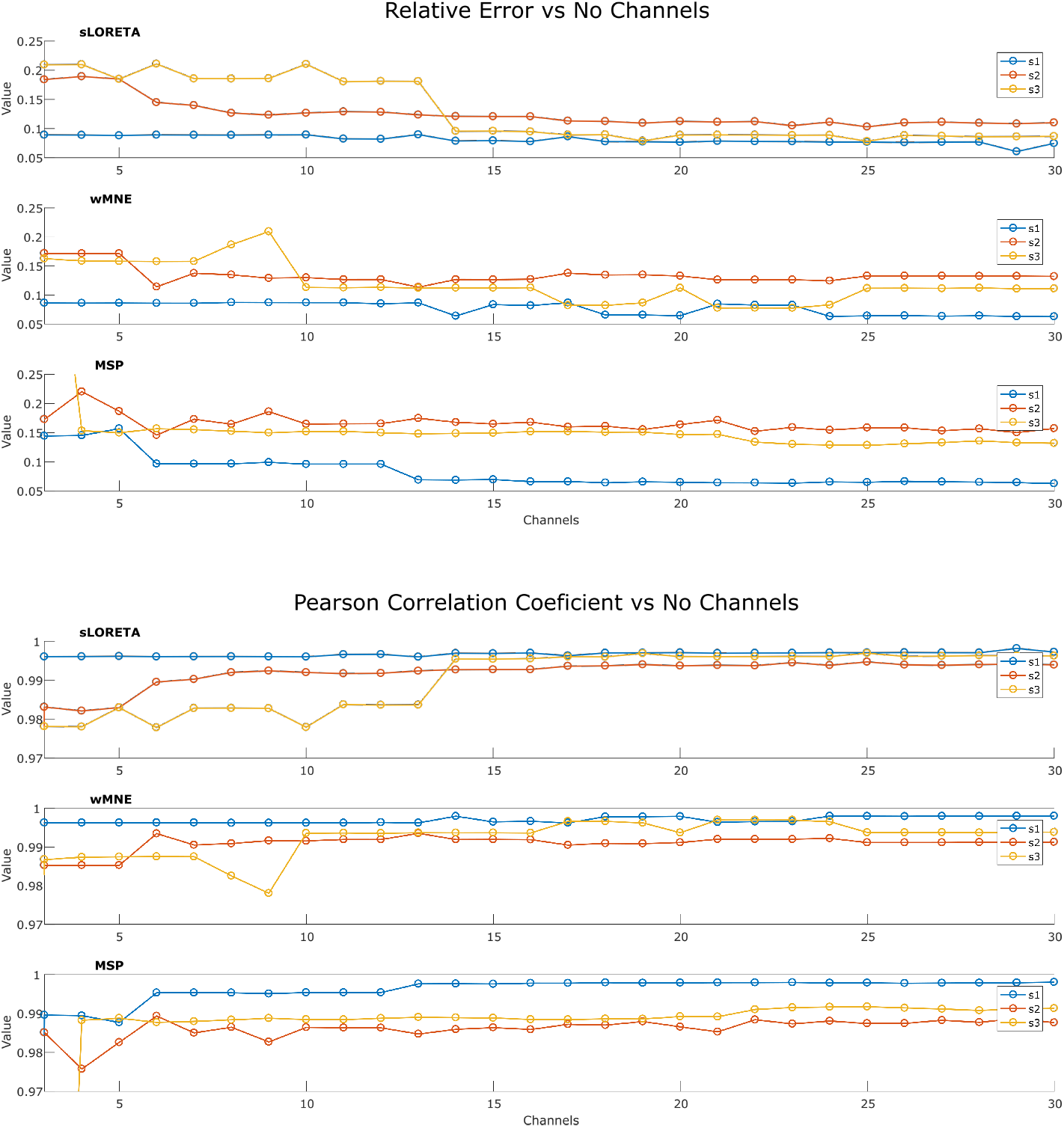
Relative error and Pearson correlation coefficient for each source, comparing the pseudo-Pareto combinations with the full set of 231 channels.

### 3.3. Localize-MI test

The ERP’s for the 61 sessions were processed by the proposed methodology with each one of the source reconstruction algorithm. During processing, the channels marked as bad channels were not taken into account neither in the optimization nor in the calculation of the values with “all channels”. The number of channels varied according to the number of channels labeled as good. The average number of good channels among the 61 sessions was 210 channels (sd = 23), in the worst case 162/256 and in the best case, 246/256 channels were considered during the optimization process. Therefore, the number of channels used varied between sessions, and the term “all channels” refers here to all channels labeled as good channels. The pseudo-Pareto front resulting from processing each one of the ERP’s and the values with all channels are presented in figure 11. It can be observed a similar tendency than in the previous tests, in which the optimized combinations obtained a lower localization error than with all the electrodes. In contrast to previous tests, there is not an exponential trend when considering the fewest number of channels, instead, a more linear behavior can be observed, where the localization error and standard deviation increases as the number of channels decreases.

**Figure 11:**
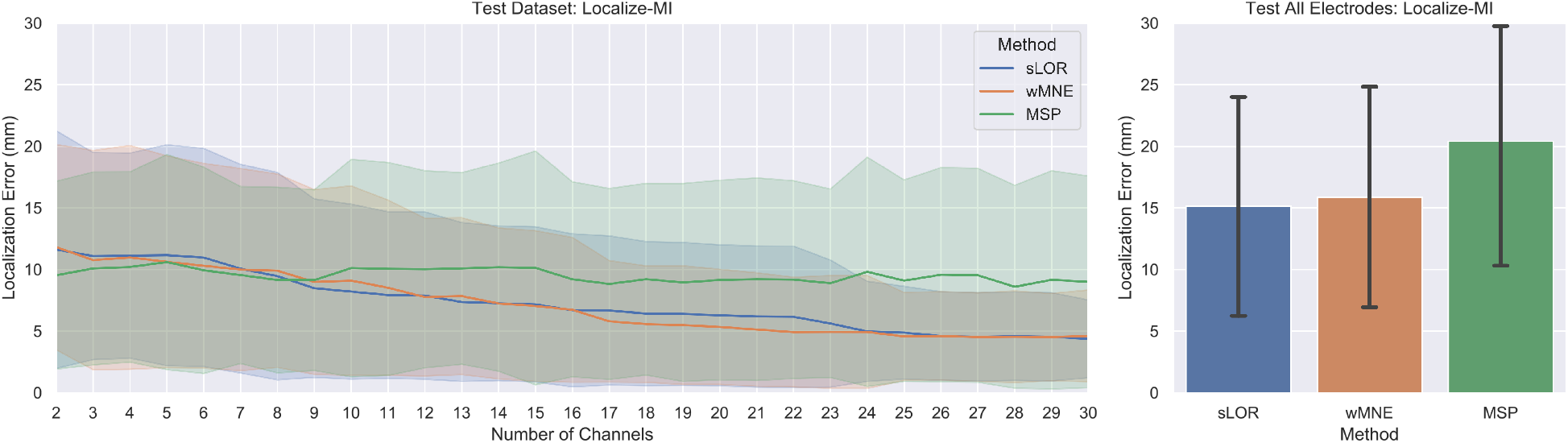
Results for the Localize-MI test. The pseudo-Pareto fronts relating the number of channels and the localization error by each source reconstruction method are presented at left. The lines represent the mean localization error across the 61 sessions and their respective colored bands represent their standard deviation. The mean localization errors across the trials and the standard deviation obtained by using all the electrodes available (all channels labeled as good) by each method are presented at right.

Table 3 presents the summary of the localization error, standard deviation and accuracy index for the Localize-MI test for the pseudo-Pareto combinations from two to sixteen channels. The accuracy index shows that more than 70% of the ERP’s were located with the same or higher accuracy than with all the electrodes when using 8 or more electrodes, and more than 60% when using four. It is noticeable that with the optimized subset of two channels, the mean localization error was lower than with all channels. In addition, the accuracy index values were 63.93%(39/61), 65.57%(40/61), and 40.98%(25/61) for sLORETA, wMNE and MSP respectively, with a marginal difference in the standard deviation for sLORETA and wMNE, when compared to the deviation obtained with all electrodes.

**Table 3.** Localize-MI test results.

| Method | Chs | Mean Loc. Error | SD | Index (%) | Method | Chs | Mean Loc. Error | SD | Index (%) | Method | Chs | Mean Loc. Error | SD | Index (%) |
| --- | --- | --- | --- | --- | --- | --- | --- | --- | --- | --- | --- | --- | --- | --- |
| sLOR | 2 | 11.62 | 9.57 | 63.93 | wMNE | 2 | 11.82 | 8.27 | 65.57 | MSP | 2 | 11.83 | 13.58 | 40.98 |
|  | 3 | 11.10 | 8.34 | 63.93 |  | 3 | 10.77 | 8.83 | 65.57 |  | 3 | 10.34 | 11.64 | 50.82 |
|  | 4 | 11.14 | 8.25 | 63.93 |  | 4 | 10.99 | 9.01 | 65.57 |  | 4 | 10.06 | 12.41 | 60.66 |
|  | 5 | 11.17 | 8.90 | 63.93 |  | 5 | 10.63 | 8.53 | 67.21 |  | 5 | 8.74 | 9.45 | 70.49 |
|  | 6 | 10.98 | 8.78 | 63.93 |  | 6 | 10.31 | 8.24 | 68.85 |  | 6 | 8.32 | 7.90 | 72.13 |
|  | 7 | 10.07 | 8.40 | 68.85 |  | 7 | 10.00 | 8.13 | 68.85 |  | 7 | 8.26 | 8.35 | 78.69 |
|  | 8 | 9.47 | 8.35 | 73.77 |  | 8 | 9.91 | 7.79 | 70.49 |  | 8 | 7.91 | 7.35 | 81.97 |
|  | 9 | 8.49 | 7.18 | 73.77 |  | 9 | 9.00 | 7.44 | 73.77 |  | 9 | 8.54 | 8.04 | 80.33 |
|  | 10 | 8.21 | 7.05 | 73.77 |  | 10 | 9.11 | 7.63 | 73.77 |  | 10 | 8.13 | 7.39 | 83.61 |
|  | 11 | 7.94 | 6.70 | 73.77 |  | 11 | 8.53 | 7.06 | 75.41 |  | 11 | 8.75 | 8.04 | 78.69 |
|  | 12 | 7.89 | 6.74 | 73.77 |  | 12 | 7.77 | 6.35 | 77.05 |  | 12 | 9.29 | 8.50 | 83.61 |
|  | 13 | 7.37 | 6.39 | 73.77 |  | 13 | 7.84 | 6.31 | 75.41 |  | 13 | 8.85 | 7.89 | 83.61 |
|  | 14 | 7.27 | 6.23 | 75.41 |  | 14 | 7.25 | 6.08 | 78.69 |  | 14 | 8.72 | 8.27 | 88.52 |
|  | 15 | 7.18 | 6.24 | 75.41 |  | 15 | 7.05 | 6.06 | 78.69 |  | 15 | 8.57 | 7.70 | 85.25 |
|  | 16 | 6.69 | 6.16 | 77.05 |  | 16 | 6.74 | 5.83 | 80.33 |  | 16 | 9.04 | 8.30 | 85.25 |
|  | All | 15.12 | 8.88 | - |  | All | 15.89 | 8.94 | - |  | All | 19.83 | 9.61 | - |

## 4. Discussion

The minimum number of electrodes required for modelling a single source is defined by the number of parameters to describe it: the localization based on three location coordinates (x,y,z) and the orientation based on three strength components in each coordinate axis (Hari and Puce (2017)). However, in distributed models based on FEM and BEM modeling, the orientation can be assumed as fixed throughout the activation period of the source, where often is assumed to be normal to the cortical surface. Therefore the number of parameters and electrodes required can be reduced. In the single-source test, the results shown in figure 6 suggest that four or five channels (where is located the inflexion point of the pesudo-Pareto’s front) are enough to estimate the source parameters with a satisfying localization accuracy when compared with a set of HD-EEG. This results are also supported by the values shown in Table 3, where the mean localization error was lower with these number of channels than when using all available electrodes.

In Sohrabpour et al. (2015) it was discussed that a plateau behavior was observed in the localization error when increasing the number of electrodes. This plateau can also be observed in the pseudo-Pareto front for all the tests performed here (figures 6, 7 and 11), where adding channels to the subset did not continue decreasing the localization error. However, the plateau did not remain flat until reaching the full number of electrodes, at some point adding channels started to increase the localization error. This effect is possible to see in the pseudo-Pareto front of the multiple-source test constrained to 60 channels, where around 23 channels, the error start increasing notably. This effect is presented because the optimization algorithm was set to identify electrode combinations with the lowest number of electrodes and the lowest localization error possible for each source; during the optimization process, the algorithm does not find the optimal for each number of electrodes, rather, it evaluates a given number and as it finds combinations with a lower number, it keeps the search in that direction. At the end, most of the combinations evaluated lied in the first third of the total electrode number. This effect was also present in the other test but for a higher number of electrodes (Not shown in the figures).

The results of the multiple-test with 231, 128, and 60 channels evidence the general effect of using a lower number of channels, where the localization error achieved the lower values when using the 231 channels in both, without optimization when using all channels, and with optimization for the subset of channels. Considering this, when constraining the search to use less electrode positions, as in the tests with 128 and 60 channels, it was possible to observe an evident increment in the localization error with and without optimization (Figure 7). This accuracy behavior supports the proposition that intermediate positions between electrodes in the 10-10 system adopted by the 10-5 electrode layout play an important role. As there are more channels, the probability of recording closer to the source of interest increases, resulting in a more accurate estimation. Moreover, the results are in line with the widely accepted concept of higher number of channels leading to a higher localization accuracy, which has been previously demonstrated and discussed (Sohrabpour et al. (2015); Song et al. (2015)).

What is new in the results is that an optimized subset of electrodes can attain similar or better localization accuracy than when using all electrodes in a HD-EEG setting (extensions of the 10-20), as shown in tables 1, 2 for a simulated dataset, and in 3 for the Localize-MI dataset. At first it can appear counter-intuitive that a reduced number of electrodes can attain such levels of accuracy. However, it should be cautiously noted that not any subset of channels can attain that accuracy. Through the combination of methods proposed in our methodology, it was possible to find combinations of channels for a particular brain source/s configuration that attained values of localization error equal or better than with HD-EEG.

Simplifying the number of electrodes by optimal selection can be beneficial for applications and systems based on sparse number of channels in medical and non-medical fields. Nowadays, there are important number of devices with EEG recording capabilities with less than 32 electrodes, and specially in the clinical setting systems with 21 electrodes based on the standard 10-20 are still the gold standard for multiple analysis (Seeck et al. (2017)). For those systems, the electrodes can be selected to estimate the activity of an area of interest; as demonstrated, the sources retrieved when using an optimized subset presented a high level of correlation with the signals obtained with the full set of electrodes (Figure 9), therefore, the source time-courses can be used for further analysis and feature extraction.

Multiple BCI systems are based on the brain activity and responses of a particular region or area of the brain e.g. classification of steady state visually evoked potential (SSVEP), or motor imaginary movements. Several works have studied the applicability, feasibility of source reconstruction in BCI classification (Ahn et al. (2012); Lindgren (2017)) and applied over motor imaginary task (Edelman et al. (2016); Li et al. (2019); Lotte et al. (2009)) reporting an increase in the classification accuracy for source-based approaches over the traditional sensor-based methods. We consider that the use of the source space has been poorly exploited in BCI systems, partially, due to the uncertainty of source reconstruction when recording with non HD-EEG systems. We believe that a selected configuration of electrodes, based on channel optimization to reconstruct the sources of specific areas, can favour the development of source-centered BCI systems with sparse number of electrodes that can surpass actual systems in accuracy, portability and comfort.

The use of low number of electrodes, inherently, reduces the preparation time, and increases the portability of the EEG systems. The implementation of the proposed methodology can increase the flexibility of brain source reconstruction in multiple studies. For example, in mobile brain/body imaging (MoBI) there is a need to make flexible and ease the acquisition of brain data (Lau-Zhu et al. (2019)), in order to perform studies in natural environments for the participants, like while exercising or working (Mehta and Parasuraman (2013)). The proposed methodology also has application in clinical settings in which is required to monitor the activity of a specific area of the brain. The electrodes required to map a particular region of the brain related to a disease can be identified with our method and placed in a patient, then, the estimated time-course can be analyzed, for example, for detecting the start of a seizure in epileptic patients, or monitor the changes in the brain activation along the time.

This optimization-based approach can enable systematic searches of electrode subsets for any given source or combination of sources that can estimate the source/s location/s. Also, the accuracy of different high-density electrode configurations widely used today can be compared for any given source scenario. In low-density systems, it can be used for evaluating the source localization quality of multiple consumer-grade EEG that proliferate nowadays Sawangjai et al. (2020); to verify to what extent they can be used, and to determine the potential brain source activity they can monitor.

The use of individual head models can be seen as a particular limitation to extend the applicability of the proposed methodology. We are aware that in multiple applications, especially the non-medical, there is no anatomical information available that allows to estimate a subject-specific model. However, in such cases, it is recommended the use of template precise models and apply a warping process using head landmarks and electrode positions (Akalin Acar and Makeig (2013)), after warping the model and co-registering with the electrodes, the forward model can be computed and used during source reconstruction.

In conclusion, the different tests performed in this study offer a multiple view of the channel optimization effects for source reconstruction in which multiple and single source cases were analyzed. Overall, the proposed methodology was able to optimize the localization error and the number of channels, by finding a subset of channels with few electrodes that attained equal or similar localization accuracy than HD-EEG. In both cases, when analyzing the Localize-MI and the synthetic EEG datasets, the sources have different localization, which allowed us to evaluate the methodology over different brain regions and source configurations. Additionally, the proposed methodology was tested on several electrode systems (10-20, 10-10, 10-5, and geodesic) and considering different head modelling methods (FEM and BEM). Independently of the variations, our proposal was able to found combinations of channels with few channels that offer equal or better localization accuracy, demonstrating the feasibility of using low-density optimized electrode combinations to localize single and multiple sources.

## 5. Data and code Availability Statement

The synthetic dataset of multiple source activity analyzed along this document can be downloaded from https://github.com/anfesogu/Ground-Truth-EEG-Dataset, where an example code for generating a synthetic dataset is also available. The forward model used for generating the synthetic EEG dataset and solving the inverse problem “New York Head model” is available at https://www.parralab.org/nyhead/ (Huang et al. (2016)). The dataset of Simultaneous human intracerebral stimulation and HD-EEG “Localize-MI” can be found in https://doi.gin.g-node.org/10.12751/g-node.1cc1ae/ (Mikulan et al. (2020)). The NSGA-II solver can be found in https://se.mathworks.com/matlabcentral/fileexchange/31166-ngpm-a-nsga-ii-program-in-matlab-v1-4 (Song (2011)).

## 6. Author contribution

AS designed and simulated all the scenarios, processed and analyzed the results and wrote the manuscript, LM designed the optimization algorithm which was adapted to source reconstruction by AS, read and wrote the manuscript, EG and MM conceived the idea of identifying optimal subsets of EEG channels for EEG source reconstruction by using optimization, discussed the scenarios to be simulated and the results, read and wrote the manuscript.

## References

Acharya, J.N., Hani, A., Cheek, J., Thirumala, P., Tsuchida, T.N., 2016. American Clinical Neurophysiology Society Guideline 2: Guidelines for Standard Electrode Position Nomenclature. Journal of Clinical Neurophysiology 33, 308–311. URL:https://journals.lww.com/clinicalneurophys/Fulltext/2016/08000/American_Clinical_Neurophysiology_Society.4.aspx, doi:10.1097/WNP.0000000000000316.

Ahn, M., Hong, J.H., Jun, S.C., 2012. Feasibility of approaches combining sensor and source features in brain–computer interface. Journal of Neuroscience Methods 204, 168–178. doi:10.1016/J.JNEUMETH.2011.11.002.

Akalin Acar, Z., Makeig, S., 2013. Effects of Forward Model Errors on EEG Source Localization. Brain Topography 2013 26:3 26, 378–396. URL: https://link.springer.com/article/10.1007/s10548-012-0274-6, doi:10.1007/S10548-012-0274-6.

Brodbeck, V., Spinelli, L., Lascano, A.M., Wissmeier, M., Vargas, M.I., Vulliemoz, S., Pollo, C., Schaller, K., Michel, C.M., Seeck, M., 2011. Electroencephalographic source imaging: A prospective study of 152 operated epileptic patients. Brain 134, 2887–2897. URL:https://academic.oup.com/brain/article/134/10/2887/324258, doi:10.1093/brain/awr243.

Chatrian, G.E., Lettich, E., Nelson, P.L., 1988a. Improved nomenclature for the ‘10%’ electrode system. American Journal of EEG Technology 28, 161–163. URL:https://www.tandfonline.com/doi/abs/10.1080/00029238.1988.11080262, doi:10.1080/00029238.1988.11080262.

Chatrian, G.E., Lettich, E., Nelson, P.L., 1988b. Modified Nomenclature for the “10%” Electrode System. Journal of Clinical Neurophysiology 5, 183–186.

Colton, D., Kress, R., 2019. Inverse Acoustic and Electromagnetic Scattering Theory. 4 ed., Springer.

Dale, A.M., Sereno, M.I., 1993. Improved localization of cortical activity by combining EEG and MEG with MRI cortical surface reconstruction: A linear approach. Journal of Cognitive Neuroscience 5, 162–176. doi:10.1162/jocn.1993.5.2.162.

Deb, K., Pratap, A., Agarwal, S., Meyarivan, T., 2002. A fast and elitist multiobjective genetic algorithm: NSGA-II. IEEE Transactions on Evolutionary Computation 6, 182–197. doi:10.1109/4235.996017.

Edelman, B.J., Baxter, B., He, B., 2016. EEG source imaging enhances the decoding of complex right-hand motor imagery tasks. IEEE Transactions on Biomedical Engineering 63, 4–14. doi:10.1109/TBME.2015.2467312.

Fischl, B., 2012. FreeSurfer. NeuroImage 62, 774–781. doi:10.1016/J.NEUROIMAGE.2012.01.021.

Friston, K., Harrison, L., Daunizeau, J., Kiebel, S., Phillips, C., Trujillo-Barreto, N., Henson, R., Flandin, G., Mattout, J., 2008. Multiple sparse priors for the M/EEG inverse problem. NeuroImage 39, 1104–1120. doi:10.1016/j.neuroimage.2007.09.048.

Fuchs, M., Wagner, M., Wischmann, H.A., 1994. Generalized minimum norm least squares reconstruction algorithmss. ISBET Newsletter 5, 8–11.

Gramfort, A., Luessi, M., Larson, E., Engemann, D.A., Strohmeier, D., Brodbeck, C., Parkkonen, L., Hämäläinen, M.S., 2014. MNE software for processing MEG and EEG data. NeuroImage 86, 446–460. doi:10.1016/J.NEUROIMAGE.2013.10.027.

Grech, R., Cassar, T., Muscat, J., Camilleri, K.P., Fabri, S.G., Zervakis, M., Xanthopoulos, P., Sakkalis, V., Vanrumste, B., 2008. Review on solving the inverse problem in EEG source analysis. Journal of NeuroEngineering and Rehabilitation 5. URL:http://www.jneuroengrehab.com/content/5/1/25, doi:10.1186/1743-0003-5-25.

Hämäläinen, M.S., Ilmoniemi, R.J., 1984. Interpreting measured magnetic fields of the brain: Estimates of current distributions.

Hämäläinen, M.S., Ilmoniemi, R.J., 1994. Interpreting magnetic fields of the brain: minimum norm estimates. Medical & Biological Engineering & Computing 32, 35–42. doi:10.1007/BF02512476.

Hari, R., Puce, A., 2017. MEG-EEG Primer. Oxford University Press, Oxford, UK. URL:https://oxfordmedicine.com/view/10.1093/med/9780190497774.001.0001/med-9780190497774-chapter-1, doi:10.1093/med/9780190497774.003.0001.

Huang, B., Buckley, B., Kechadi, T.M., 2010. Multi-objective feature selection by using NSGA-II for customer churn prediction in telecommunications. Expert Systems with Applications 37, 3638–3646. doi:10.1016/j.eswa.2009.10.027.

Huang, Y., Parra, L.C., Haufe, S., 2016. The New York Head—A precise standardized volume conductor model for EEG source localization and tES targeting. NeuroImage 140, 150–162. doi:10.1016/j.neuroimage.2015.12.019.

Iwaki, S., Ueno, S., 1998. Weighted minimum-norm source estimation of magnetoencephalography utilizing the temporal information of the measured data. Journal of Applied Physics 83, 6441. URL:https://aip.scitation.org/doi/abs/10.1063/1.367732, doi:10.1063/1.367732.

Jasper, H., 1958. The ten-twenty electrode system of the international federation. Electroencephalography and Clinical Neurophysiology 10, 370–375. URL:https://ci.nii.ac.jp/naid/10017996828.

Jatoi, M.A., Kamel, N., 2018. Brain source localization using reduced EEG sensors. Signal, Image and Video Processing 12, 1447–1454. doi:10.1007/s11760-018-1298-5.

Jurcak, V., Tsuzuki, D., Dan, I., 2007. 10/20, 10/10, and 10/5 systems revisited: Their validity as relative head-surface-based positioning systems. NeuroImage 34, 1600–1611. doi:10.1016/j.neuroimage.2006.09.024.

Kalyanmoy Deb, 2001. Multi-Objective Optimization using Evolutionary Algorithms | Wiley. volume 16. John Wiley & Sons.

Kee, C.Y., Ponnambalam, S.G., Loo, C.K., 2015. Multi-objective genetic algorithm as channel selection method for P300 and motor imagery data set. Neurocomputing 161, 120–131. doi:10.1016/J.NEUCOM.2015.02.057.

Lantz, G., Grave de Peralta, R., Spinelli, L., Seeck, M., Michel, C.M., 2003. Epileptic source localization with high density EEG: How many electrodes are needed? Clinical Neurophysiology 114, 63–69. doi:10.1016/S1388-2457(02)00337-1.

Lau-Zhu, A., Lau, M.P., McLoughlin, G., 2019. Mobile EEG in research on neurodevelopmental disorders: Opportunities and challenges. Developmental Cognitive Neuroscience 36, 100635. doi:10.1016/J.DCN.2019.100635.

Li, M.A., Wang, Y.F., Jia, S.M., Sun, Y.J., Yang, J.F., 2019. Decoding of motor imagery EEG based on brain source estimation. Neurocomputing 339, 182–193. doi:10.1016/J.NEUCOM.2019.02.006.

Lindgren, J.T., 2017. As above, so below? Towards understanding inverse models in BCI. Journal of Neural Engineering 15, 012001. URL:https://iopscience.iop.org/article/10.1088/1741-2552/aa86d0 https://iopscience.iop.org/article/10.1088/1741-2552/aa86d0/meta, doi:10.1088/1741-2552/AA86D0.

López, J.D., Litvak, V., Espinosa, J.J., Friston, K., Barnes, G.R., 2014. Algorithmic procedures for Bayesian MEG/EEG source reconstruction in SPM. NeuroImage 84, 476–487. doi:10.1016/j.neuroimage.2013.09.002.

Lotte, F., Lécuyer, A., Arnaldi, B., 2009. FuRIA: An inverse solution based feature extraction algorithm using fuzzy set theory for brain-computer interfaces. IEEE Transactions on Signal Processing 57, 3253–3263. doi:10.1109/TSP.2009.2020752.

Mehta, R.K., Parasuraman, R., 2013. Neuroergonomics: a review of applications to physical and cognitive work. Frontiers in Human Neuroscience 0, 889. doi:10.3389/FNHUM.2013.00889.

Mikulan, E., Russo, S., Parmigiani, S., Sarasso, S., Zauli, F.M., Rubino, A., Avanzini, P., Cattani, A., Sorrentino, A., Gibbs, S., Cardinale, F., Sartori, I., Nobili, L., Massimini, M., Pigorini, A., 2020. Simultaneous human intracerebral stimulation and HD-EEG, ground-truth for source localization methods. Scientific Data 7, 1–8. URL:www.nature.com/scientificdata, doi:10.1038/s41597-020-0467-x.

Moctezuma, L.A., Molinas, M., 2020a. EEG Channel-Selection Method for Epileptic-Seizure Classification Based on Multi-Objective Optimization. Frontiers in Neuroscience 14, 593. URL:www.frontiersin.org, doi:10.3389/fnins.2020.00593.

Moctezuma, L.A., Molinas, M., 2020b. Multi-objective optimization for EEG channel selection and accurate intruder detection in an EEG-based subject identification system. Scientific Reports 10, 1–12. URL:https://doi.org/10.1038/s41598-020-62712-6, doi:10.1038/s41598-020-62712-6.

Oostenveld, R., Praamstra, P., 2001. The five percent electrode system for high-resolution EEG and ERP measurements. Clinical Neurophysiology 112, 713–719. doi:10.1016/S1388-2457(00)00527-7.

Pascual-Marqui, R., 1999. Review of Methods for Solving the EEG Inverse Problem. International Journal of Bioelectromagnetism 1, 75–86.

Pascual-Marqui, R.D., 2002. Standardized low-resolution brain electromagnetic tomography (sLORETA): technical details. Methods and findings in experimental and clinical pharmacology 24 Suppl D, 5–12.

Sawangjai, P., Hompoonsup, S., Leelaarporn, P., Kongwudhikunakorn, S., Wilaiprasitporn, T., 2020. Consumer Grade EEG Measuring Sensors as Research Tools: A Review. IEEE Sensors Journal 20, 3996–4024. doi:10.1109/JSEN.2019.2962874.

Seeck, M., Koessler, L., Bast, T., Leijten, F., Michel, C., Baumgartner, C., He, B., Beniczky, S., 2017. The standardized EEG electrode array of the IFCN. doi:10.1016/j.clinph.2017.06.254.

Sinha, S.R., Sullivan, L., Sabau, D., San-Juan, D., Dombrowski, K.E., Halford, J.J., Hani, A.J., Drislane, F.W., Stecker, M.M., 2016. American Clinical Neurophysiology Society Guideline 1: Minimum Technical Requirements for Performing Clinical Electroencephalography. Journal of Clinical Neurophysiology 33, 303–307. URL:https://journals.lww.com/clinicalneurophys/Fulltext/2016/08000/American_Clinical_Neurophysiology_Society.3.aspx, doi:10.1097/WNP.0000000000000308.

Själander, M., Jahre, M., Tufte, G., Reissmann, N., 2019. {EPIC}: An Energy-Efficient, High-Performance {GPGPU} Computing Research Infrastructure.

Sohrabpour, A., Lu, Y., Kankirawatana, P., Blount, J., Kim, H., He, B., 2015. Effect of EEG electrode number on epileptic source localization in pediatric patients. Clinical Neurophysiology 126, 472–480. doi:10.1016/j.clinph.2014.05.038.

Soler, A., Giraldo, E., Molinas, M., 2020a. Low-density EEG for Source Activity Reconstruction using Partial Brain Models, in: Proceedings of the 13th International Joint Conference on Biomedical Engineering Systems and Technologies, SCITEPRESS - Science and Technology Publications. pp. 54–63. URL: http://www.scitepress.org/DigitalLibrary/Link.aspx?doi=10.5220/0008972500540063, doi:10.5220/0008972500540063.

Soler, A., Muñoz-Gutiérrez, P.A., Bueno-López, M., Giraldo, E., Molinas, M., 2020b. Low-Density EEG for Neural Activity Reconstruction Using Multivariate Empirical Mode Decomposition. Frontiers in Neuroscience 14, 175. doi:10.3389/fnins.2020.00175.

Song, J., Davey, C., Poulsen, C., Luu, P., Turovets, S., Anderson, E., Li, K., Tucker, D., 2015. EEG source localization: Sensor density and head surface coverage. Journal of Neuroscience Methods 256, 9–21. doi:10.1016/j.jneumeth.2015.08.015.

Song, L., 2011. NGPM a NSGA-II Program in Matlab v1.4. MATLAB Central File Exchange URL:https://www.mathworks.com/matlabcentral/fileexchange/31166-ngpm-a-nsga-ii-program-in-matlab-v1-4.

Srinivas, N., Deb, K., 1994. Muiltiobjective Optimization Using Nondominated Sorting in Genetic Algorithms. Evolutionary Computation 2, 221–248. doi:10.1162/evco.1994.2.3.221.

Stoyell, S.M., Wilmskoetter, J., Dobrota, M.A., Chinappen, D.M., Bonilha, L., Mintz, M., Brinkmann, B.H., Herman, S.T., Peters, J.M., Vulliemoz, S., Seeck, M., Hämäläinen, M.S., Chu, C.J., 2021. High-Density EEG in Current Clinical Practice and Opportunities for the Future. Journal of clinical neurophysiology : official publication of the American Electroencephalographic Society 38, 112–123. URL:https://journals.lww.com/clinicalneurophys/Fulltext/2021/03000/High_Density_EEG_in_Current_Clinical_Practice_and.6.aspx, doi:10.1097/WNP.0000000000000807.

Suarez, E., Viegas, M.D., Adjouadi, M., Barreto, A., 2000. Relating induced changes in EEG signals to orientation of visual stimuli using the ESI-256 machine, in: Biomedical Sciences Instrumentation, Instrument Society of America. pp. 33–38. URL:https://europepmc.org/article/med/10834205.

Tucker, D.M., 1993. Spatial sampling of head electrical fields: the geodesic sensor net. Electroencephalography and Clinical Neurophysiology 87, 154–163. doi:10.1016/0013-4694(93)90121-B.

Vorwerk, J., Cho, J.H., Rampp, S., Hamer, H., Knösche, T.R., Wolters, C.H., 2014. A guideline for head volume conductor modeling in EEG and MEG. NeuroImage 100, 590–607. doi:10.1016/j.neuroimage.2014.06.040.

